# Enhanced ISGylation reduces respiratory distress following *Francisella novicida* infection

**DOI:** 10.1101/2023.09.19.558558

**Authors:** Ellen M. Upton, Emma K. Luhmann, Yifeng Zhang, Brittany M. Ripley, David K. Meyerholz, Lilliana Radoshevich

## Abstract

The Interferon-Stimulated Gene 15 (ISG15) is a ubiquitin-like protein induced by viral and bacterial infection. ISG15 covalently modifies host and pathogenic proteins in a process called ISGylation. Yet, the consequences of ISGylation on protein fate and function remain to be determined. Here we sought to assess whether ISGylation would be protective following bacterial pneumonia caused by *Francisella novicida.* We found that infection with *F. novicida* induces ISGylation both *in vitro* in macrophages and *in vivo* in the lung, liver, and spleen of mice infected intranasally. Surprisingly, ISG15 and ISGylation do not affect bacterial burden in the lung *in vivo*, but in a model of enhanced ISGylation (*usp18^C61A/C61A^*) mice have decreased respiratory distress relative to *Isg15^-/-^* animals. In order to understand the mechanism which underlies this phenotype, we mapped the ISGylome of *F. novicida-*infected mouse lungs using label-free quantitative mass spectrometry and identified enrichment in ISGylation of proteins involved in the innate immune response and cytosolic nucleotide signaling. We validated ISGylation of the sterile alpha motif and HD-containing protein 1 (SAMHD1) via immunoprecipitation. SAMHD1 depletes cytosolic dinucleotide stores critical for retroviral replication but it is unknown how its activity could affect bacterial infection. Structure-function analysis indicates that ISG15 modification sites in *usp18^C61A/C61A^* mice could prevent SAMHD1 dimerization and therefore abrogate function. Accordingly, deletion of SAMHD1 in fibroblasts with enhanced ISGylation reduces bacterial load. Taken together, unchecked ISGylation plays a protective role in *F. novicida* infection in vivo through improved respiratory function. Thus, inhibiting USP18 may be a promising therapeutic strategy for both viral and bacterial pneumonia.

**Author summary:** *Francisella tularensis* is a bacterial pathogen responsible for the disease tularemia, which can result in severe respiratory infection if as few as ten bacteria are inhaled. Our cells have many ways of managing infections, including the production of proteins designed to fight off foreign pathogens. One protein produced following infection is the interferon-stimulated gene 15 (ISG15). ISG15 is a ubiquitin-like molecule, meaning that it can be chemically attached to other proteins. When bound ISG15 changes the stability, interacting partners, or function of its target in a process termed ISGylation. Here we show that ISG15 is produced following infection with *Francisella.* We found that enhanced ISGylation led to less severe respiratory symptoms. To better understand the mechanism by which ISGylation protects from infection we identified the ISG15-modified proteins in the lung using mass-spectrometry-based proteomics. We found protein targets that are involved in the control of immune signaling pathways including sterile alpha motif and HD-containing protein 1 (SAMHD1) which, when deleted in cells with enhanced ISGylation, leads to better bacterial clearance. Together, we show that enhanced ISGylation plays a protective role following bacterial pneumonia, indicating that targeting this pathway could prove a beneficial therapeutic in both bacterial and viral respiratory diseases.

## Introduction

*Francisella tularensis* is a Gram-negative bacterial pathogen that is the etiological agent of tularemia. *Francisella* can be ingested from contaminated food or water, vector-borne which causes ulcerative lesions in the skin, or aerosolized leading to respiratory infection and severe pneumonia [1, 2]. Since fewer than ten colony-forming units (CFUs) of *F. tularensis* are sufficient to cause disease and infected individuals present with rapid and life-threatening pneumonia, *F. tularensis* is classified as a Tier 1 select agent [2-4]. There is currently no vaccine available; however, there is an attenuated live vaccine strain (LVS) and a related-environmental strain *F. tularensis* subsp. *novicida (*also called *F. novicida)* which are commonly used in Biosafety Level 2 (BSL2) research studies [1, 3, 5-8]*. F. novicida* infects mice at a low multiplicity of infection and disease progression in this model can recapitulate many aspects of infection with more virulent strains of *F. tularensis* [5, 7, 9, 10].

Once the bacteria reach the lower respiratory tract, they infect alveolar macrophages and type II alveolar epithelial cells, rapidly replicate, and spread through macrophages to secondary sites of infection [2, 11-15]. *Francisella* evades the initial activation of host defense responses due to its unique lipopolysaccharide makeup that is not recognized by TLR4 [16, 17]. Upon entry into professional phagocytes, *Francisella* thwarts the oxidative burst and disrupts phagosomal integrity to escape into the cytosol and ensure its intracellular survival in macrophages and neutrophils [18-20]. Intracellular *Francisella* activates the AIM2 inflammasome when bacterial DNA is released into the cytosol during infection [21]. AIM2 is an Interferon-induced sensor, however, deletion of the interferon receptor (IFNAR^-/-^) leads to increased protection against *Francisella species* rather than susceptibility [15, 21-25]. Type I Interferon upregulates hundreds of Interferon-Stimulated Genes (ISGs), some of which have been well characterized in viral infections but not in the context of *Francisella* infection.

One such protein, ISG15, is induced by bacterial DNA in the cytosol following *Listeria monocytogenes* infection and acts as an antibacterial effector when properly regulated [26]. ISG15 is a ubiquitin-like molecule which conjugates to protein substrates following activation by an E1 enzyme UBE1L, conjugation by an E2 enzyme UBCH8, and ligation to protein substrates by three known E3 enzymes (HERC5/6, HHARI, and TRIM25) [27-30]. The process is reversed by the ISG15-decongugase USP18, which is also Interferon-induced [27, 31, 32]. Additionally, ISG15 functions as a cytokine in response to infection and can activate increased production of Interferon-γ [33-39]. ISG15 covalently modifies both host and viral proteins in a process called ISGylation, which inhibits viral replication and budding [33, 40-45]. More recent work suggests that ISG15 can also act as an antibacterial effector [26, 46-48]. However, the specific mechanism of action and consequences of ISG15 modification on host proteins following infection have yet to be fully explored.

Here we sought to assess whether ISG15 functions as an antibacterial effector following respiratory infection with *F. novicida*. We found that ISG15 and ISGylation are significantly induced following *F. novicida* infection both *in vitro* and *in vivo*. We leveraged models of absent (*isg15^-/-^*) and enhanced ISGylation (*usp18^C61A/C61A^*) to determine how the presence and regulation of ISG15 impacts bacterial burden and observed no significant differences in bacterial load following infection *in vivo*. However, enhanced ISGylation results in improved airway function. We further mapped the complete ISGylome of *F. novicida-*infected lungs via quantitative, label-free proteomics to assess potential mechanisms of reduced pathology in the enhanced ISGylation model. We identified an enrichment of ISGylated proteins involved in innate immune signaling in the lungs of *usp18^C61A/C61A^* mice and were able to validate the sterile alpha motif and histidine-aspartate-domain-containing protein 1 (SAMHD1) as a ISG15 substrate. Finally, we observed that SAMHD1-deficiency in *usp18^C61A/C61A^* fibroblasts and THP-1 derived macrophages restricts bacterial burden.

## Results

### ISG15 and ISGylation are induced by *Francisella novicida* infection

A growing body of literature demonstrates the importance of ISG15 in the host response to bacterial infections [26, 46-48]. Our primary goal in this study was to assess whether inhibition of USP18, which leads to enhanced ISGylation, could be protective in bacterial lung infections. Therefore, we first assessed whether ISG15 and ISGylation are induced following infection with the intracellular pathogen *Francisella novicida.* We utilized bone marrow-derived macrophages (BMDMs) to model infection of alveolar macrophages. We observed robust induction of ISG15 and ISGylation following infection with *F. novicida* in these cells (Fig 1A). Additionally, a different pattern of slower migrating bands, indicating proteins covalently modified by ISG15, emerged in wild-type BMDMs relative to those obtained from *usp18^C61A/C61A^* mice. We wanted to determine if ISG15 and ISGylation convey a protective effect, as is the case for other bacterial infections [26, 34, 46-49]. We challenged BMDMs with *F. novicida* at an MOI of 10 and surprisingly observed a significant increase of colony forming units (CFUs) in the *usp18^C61A/C61A^*-derived macrophages (Fig 1B) relative to both wildtype and *isg15^-/-^* macrophages. This suggests that too much ISGylation alters the macrophage response to infection tipping the balance in favor of *Francisella* replication at twenty hours post infection *in vitro*. In order to determine whether enhanced ISGylation would be protective or deleterious in the complex tissue environment of the lung, we subsequently tested the consequences of enhanced or absent ISGylation *in vivo*.

**Fig 1.**
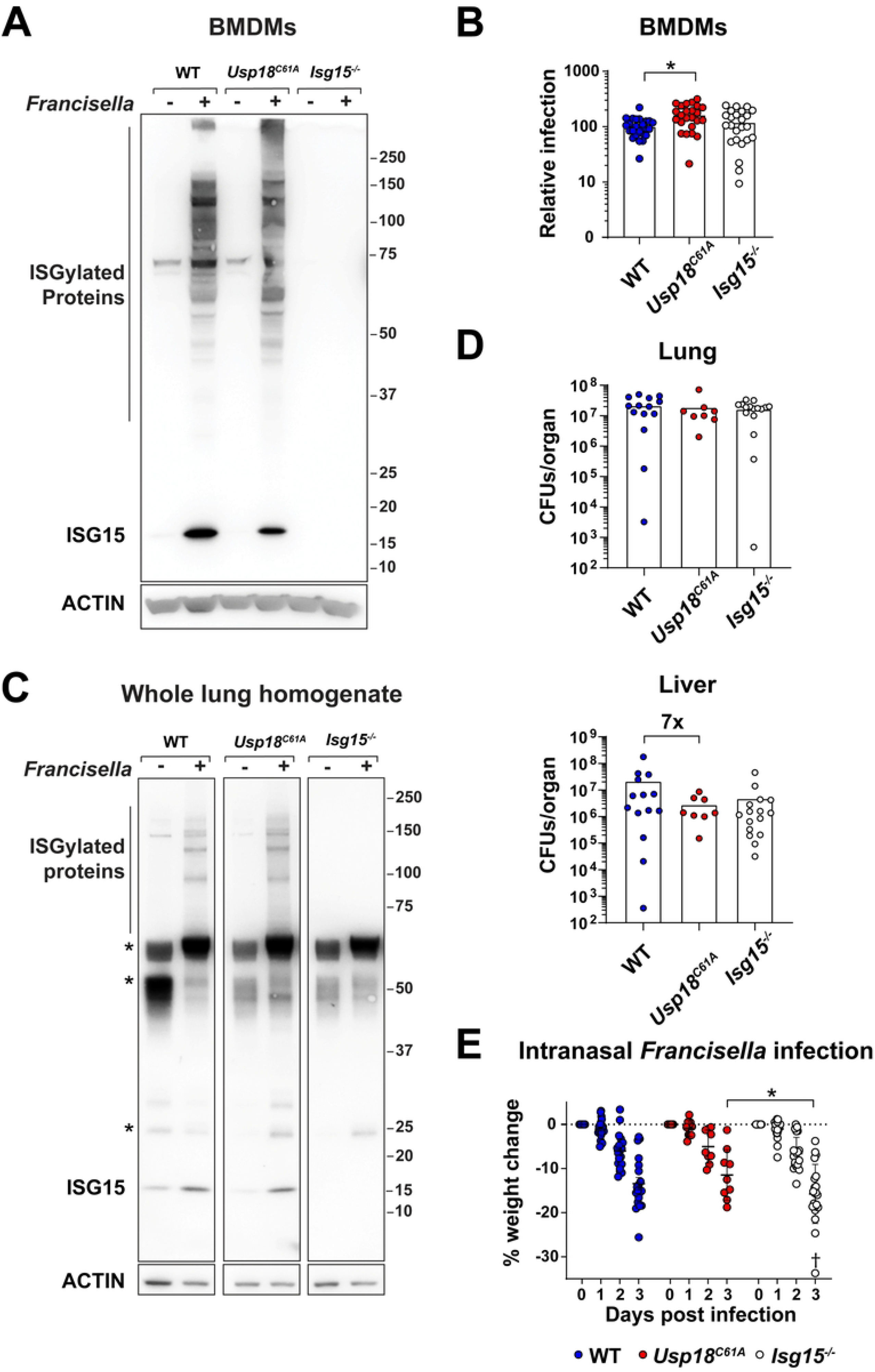
Induction and effect of ISG15 following *F. novicida* infection. (A) SDS-PAGE analysis of lysate from BMDMs infected with *F. novicida* for 24, probed for ISG15 and actin. (B) CFUs from BMDMs infected with *F. novicida,* MOI of 10 for 20 hours. Wild-type values were normalized to one hundred; data from three independent experiments with n=6/group is shown analyzed by Kruskal-Wallis test with Dunn’s multiple comparisons, adjusted p-value is shown for BMDMs *p=0.0101. (C-E) Mice were infected with 100 CFUs of *F. novicida* intranasally and monitored for 48 or 72 hours after which they were sacrificed, and organs were homogenized in PBS. (C) SDS-PAGE of lung homogenate probed for ISG15 and actin following 48-hour infection, *’s indicate background bands. (D) CFUs were enumerated from homogenized organs, data from two independent experiments are shown with n=4-9 mice/group. (E) Percent weight change from day zero was calculated and graphed for *Francisella-*infected mice over the course of infection, mice were monitored daily and sacrificed on day three or before if ≥20% of body weight was lost. † represents specimen found dead in pen (DIP). Compiled data from two independent experiments are shown n=4-9 mice/group, analyzed by two-way ANOVA with Tukey’s test post hoc, adjusted p-value is shown *p=0.0142.

*F. novicida* first infects alveolar macrophages [50] and epithelial cells *in vivo* [11]. The bacteria subsequently replicate within invading neutrophils [20], inhibiting apoptosis of PMNs [20, 51] long enough to spread to other organs through the Trojan horse mechanism via circulating macrophages and monocytes that phagocytose neutrophils [52]. We employed an intranasal infection strategy to replicate the severe respiratory form of the disease in mice with enhanced or absent ISGylation. As we observed *in vitro*, ISG15 and ISGylation were induced in the lung (Fig 1C), liver, and spleen (S1 Fig A-B) following intranasal delivery of *F. novicida*. These data indicate that ISG15 induction contributes to the host response during *F. novicida* infection. We next sought to determine the physiological relevance of ISGylation to host defense from this pathogen. We initially quantified bacterial load in the lung, liver, and spleen at either 48 or 72 hours post-infection (Fig 1D and S1 Fig C-D). Unlike the *in vitro* data, we did not see a statistically significant difference in bacterial quantity between the genotypes at either time point in the lung. There was, however, a seven-fold decrease in bacteria which had disseminated from the lung to the liver in conditions of enhanced ISGylation (Fig 1D). While the decrease in spread of bacteria to the liver was not statistically significant, we next assessed if it was physiologically relevant to host defense in infected mice. Interestingly, when observing mouse weights and behavior over time following infection, the ISG15-deficient animals lost significantly (p=0.0142) more weight with generally poorer body condition scores than *usp18^C61A/C61A^* mice (Fig 1E). *Francisella* replicates rapidly up to millions of colony-forming units per organ before innate immune detection results in extensive immune infiltrate [53], occluding normal respiration [9, 54, 55]. Since we found that *usp18^C61A/C61A^* macrophages could not control initial intracellular bacterial replication as well as wildtype cells *in vitro* but mice appeared phenotypically resistant to infection *in vivo*, we hypothesize that lung function may be protected by enhanced ISGylation.

### Enhanced ISGylation protects respiratory function following infection

We sought to test our hypothesis that ISG15 modification affects lung pathology of *F. novicida* infected animals by examining lung histology from mice with wildtype, absent ISG15, or enhanced ISGylation three days post infection. We examined sections of infected lung and saw a wide range of lesion severity within genotypes, which correlates with the inherent variability of infection that we and others [56] have observed in colony forming units between animals infected with *Francisella* via the intranasal route (Fig 2A). Despite this variability, the *usp18^C61A/C61A^* mice had fewer regions of necrotizing inflammation than ISG15-deficient or wildtype mice indicating less severe disease (Fig 2A). Examination of tissue sections can be limited by the sectioning plane through the organ. Due to the wide range of pathological phenotypes observed, we sought out a quantitative measure of lung airway function. Following intranasal *F. novicida* infection, we used a whole-body plethysmograph to measure daily changes in respiration following infection (Fig 2B, 2C). We found that 72 hours post-infection there was a distinct and significant difference (p=0.0248) between the respiration rates and breathing patterns of *usp18^C61A/C61A^* mice and *isg15^-/-^* mice (Fig 2C). PenH (plethysmography and increases in enhanced pause) is a published measure that correlates with airway restriction in studies of allergy and asthma and allows for repeated non-invasive measurements of lung capacity [57]. Using this measure, airway resistance was increased in *isg15^-/-^* mice on Day 3 post-infection relative to *usp18^C61A/C61A^* mice (Fig 2D). Notably, this indicates a protective effect of ISGylation in controlling airway restriction, which compares to the relative cellular infiltration seen at histological analysis. At the same time, we also collected data on breaths per minute. Compared to the depressed respiratory rate seen in WT and ISG15, the *usp18^C61A/C61A^* mice had a significantly higher respiratory rate (p=0.0194, p=0.0076) that was closer to the baseline values at the start of the experiment (Fig 2E). This suggests that enhanced ISGylation initially favors bacterial replication in macrophages *in vitro* but is ultimately clinically protective during bacterial pneumonia *in vivo*.

**Fig 2.**
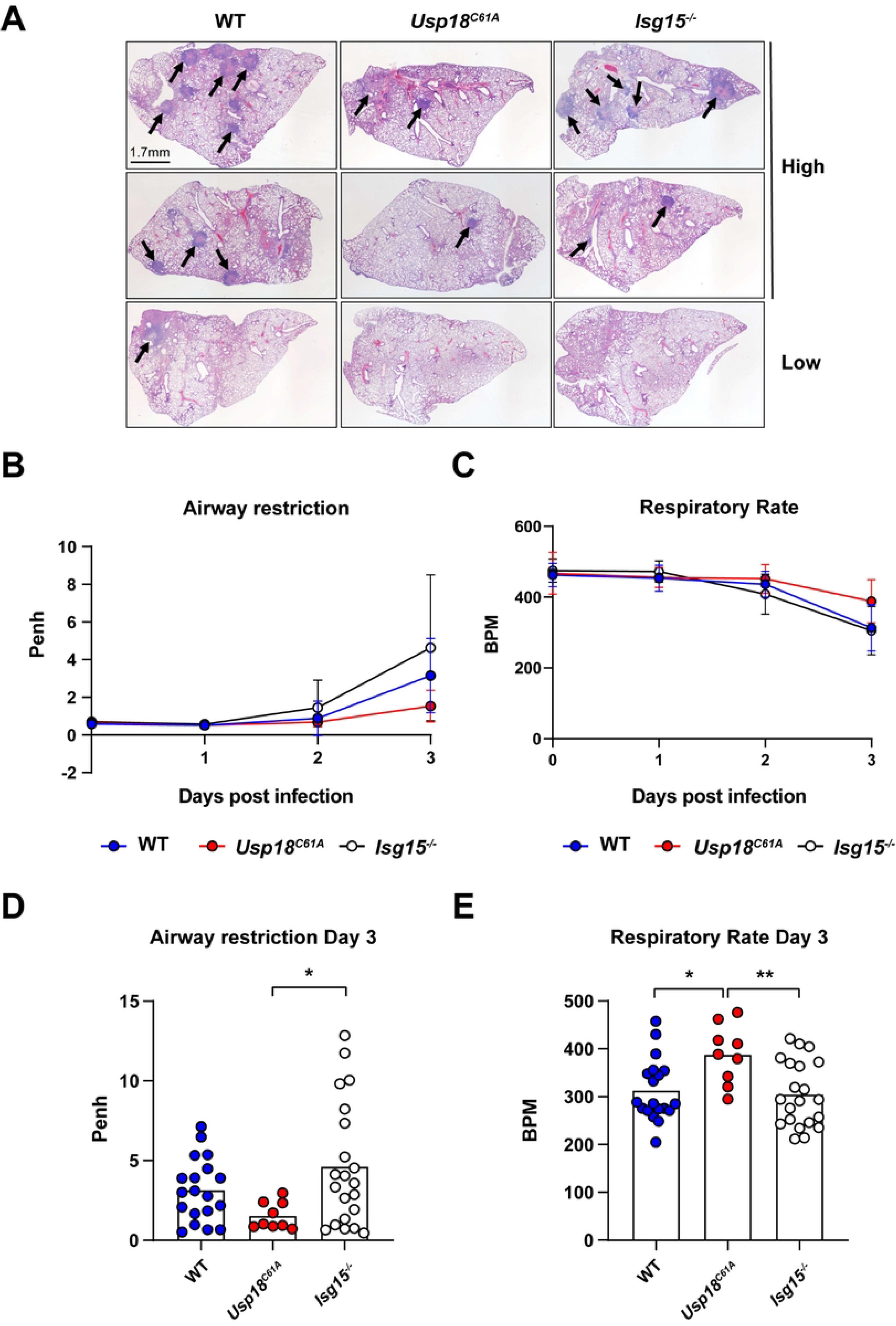
Hyper-ISGylation results in decreased airway restriction in the lungs of *Francisella-*infected mice. Mice were infected intranasally with 100 CFUs of *F. novicida* on day 0. (A) Representative H&E figures from the lungs of mice on day three post-infection distinguished between mice that had more severe disease (high) and less severe disease (low). Arrows denote areas of necrotizing pyogranulomatous inflammation. (B-E) Mice were monitored daily for airway restriction and respiration rate using a whole-body plethysmograph, combined data from 2-3 individual experiments with n= 4-9 mice/group is shown. (B-C) Show values of Penh and respiration rate over the course of infection. (D-E) Show values of Penh and respiratory rate on day three post-infection respectively were plotted. Significance was determined via an ordinary one-way ANOVA with Tukey’s test post hoc, Adjusted p-values are shown; (D) *p=0.0248, (E) *p=0.0194, **p=0.0076

### Lung ISGylome following acute *F. novicida* infection reveals a subset of USP18-regulated targets

Since phenotypic differences in histology and lung function were observed due to dysregulated ISGylation, we sought to map ISG15-modified substrates in the lungs of *F. novicida-*infected mice. We hypothesized that specific substrates present in *usp18^C61A/C61A^* mice but absent in wildtype mice could underlie the mechanism by which lung function may be altered. Our previous work established a method to simultaneously and quantitatively map the ubiquitylome and ISGylome *in vivo* following *Listeria monocytogenes* infection in the liver [49]. While others use USP18 deletion [58] or have proposed to leverage ISG15-directed viral proteases to identify ISG15 sites [59], we make use of ISG15-deficient cells or mice. In so doing, we can identify and quantify relative levels of peptides with diglycine-modified lysine adducts, which are indicative of ISGylation or ubiquitylation sites, prior to and following infection. We used individual mice as biological replicates and leveraged label-free quantitative proteomics using the open-source software MSFragger [60, 61] and Perseus [62] to analyze the data. We applied this method to bulk tissue from the lungs of mice either infected with *Francisella novicida* for 72 hours or non-infected controls. *Isg15^-/-^* mice permit us to map the strict ubiquitylome, which we then compare to wildtype to differentiate between ISG15 and ubiquitin sites. Using four biological replicates per genotype from uninfected and infected mice, we identified 5871 total sites across twenty-four samples, including sites modified by several PTMs. We consider a site quantified if identified in a minimum of three biological replicates across all groups. Using these criteria, we quantified 3003 total sites. Of the quantified sites we find that 1512 are significantly regulated either by genotype, infection, or their intersection using two-way ANOVA (S1 Table). Following unsupervised hierarchical clustering biological replicates of infected wildtype, ISG15-deficient, and *usp18^C61A/C61A^* mice clustered based on genotype and infection meaning that ISGylation following infection dramatically affects the pattern and quantity of diglycine sites (Fig 3A). Uninfected genotypes, however, did not distinctly differ from each other which highlights that prior to infection in the lung, the majority of diglycine sites are not regulated by ISG15 expression levels or the lack of ISG15. These sites are also present in the ISG15-deficient mice which indicates they are likely ubiquitin sites (Fig 3A). This is consistent with our previously published data in the liver. We considered 574 sites to be bone fide ISGylation sites due to their absence in our *isg15^-/-^* mouse samples. These sites consisted of three groups: Clusters 2, 4, and 5 (Fig 3A). Cluster 5 sites are present in wildtype and *usp18^C61A/C61A^* mice and consists of 322 sites on 138 proteins. Whereas Cluster 2 is enriched in wildtype, but absent or reduced in *usp^18C61A/C61A^* mice; under these conditions we identified 131 peptides on 98 proteins. Cluster 4 is solely present in *usp18^C61A/C61A^* conditions and includes 121 modified peptides on 88 proteins. The majority of sites were ubiquitin modifications from Cluster 3, corresponding to 831 peptides across 404 proteins which were induced by infection (S1 Table). Interestingly, Cluster 1 sites appear to be ubiquitinated prior to infection, though some sites were variably lost across genotypes. This could indicate a one-to-one replacement; however our method cannot distinguish between ubiquitin and ISG15 under these conditions. Overall, nearly 19% of quantified diglycine peptide adducts are ISG15 sites, which is similar to what we observed following *Listeria* infection of the liver [49]. Unlike the *Listeria* infected liver, however, where infection induced a loss of ubiquitin sites, *Francisella* provokes a gain of both ISG15 and ubiquitin sites. Whether this is due to unique properties of each pathogen, or the tissue-specific response of each organ remains to be determined.

**Fig 3.**
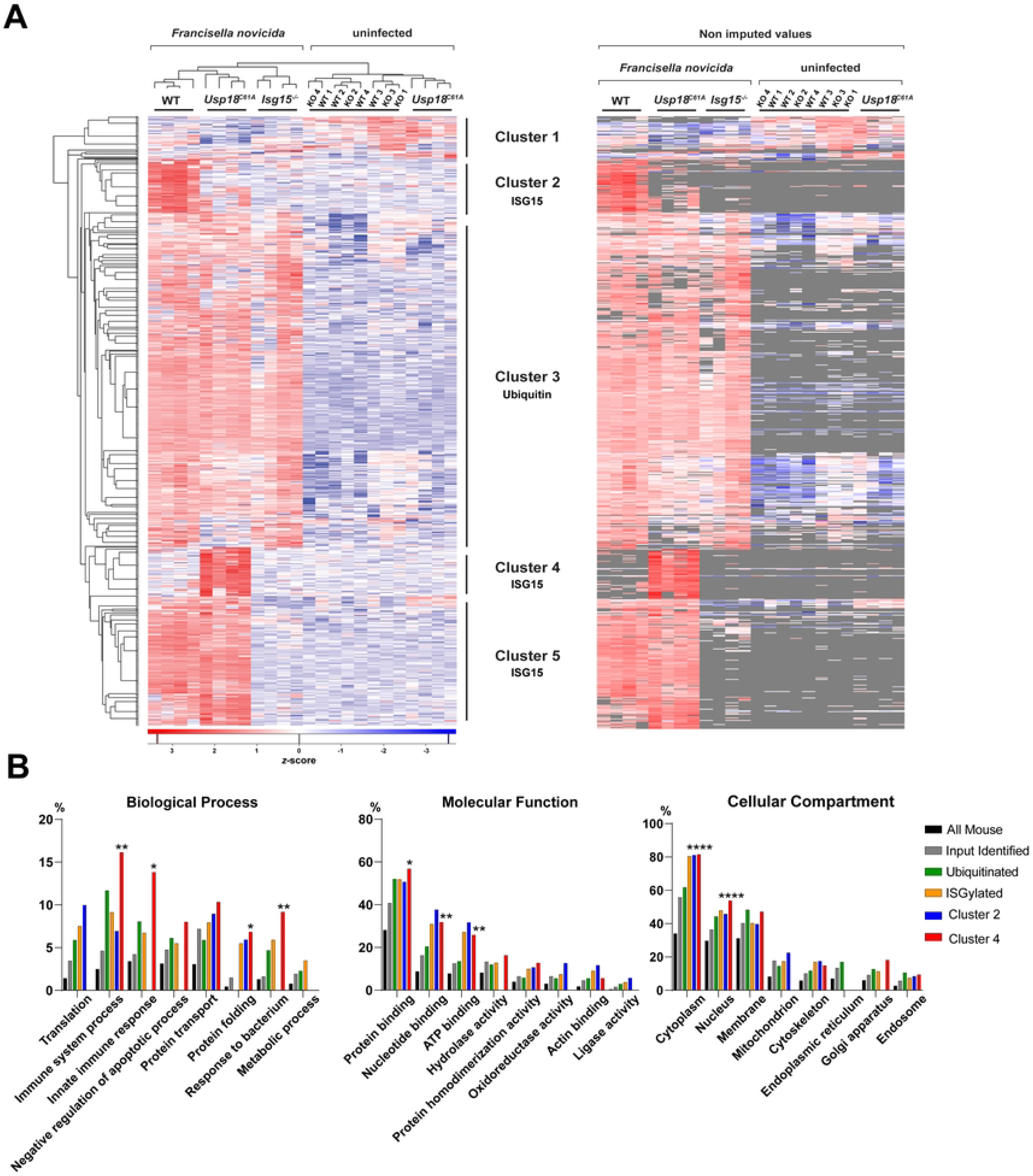
The ISGylome of *Francisella*-infected lungs highlights differentially regulated ISG15 modifications in hyper-ISGylation conditions. (A) Heatmap showing significantly enriched GlyGly(K) sites from a two-way ANOVA after non-supervised hierarchical clustering. On the right side, the heatmap is shown with missing values in gray. Five major clusters can be observed corresponding to ISG15 sites (clusters 2, 4, and 5) and ubiquitin sites (clusters 1 and 3) (B) GO analysis of Ubiquitinated peptides enhanced during infection (Cluster 3), ISGylated peptides (Cluster 2,4, and 5), and the individual Cluster 2 and Cluster 5 enhanced in wild-type or *usp18^C61A/C61A^* samples respectively were assessed relative to all proteins identified the lung samples (gray) and all mouse proteins in the Uniprot/Swiss-Prot database (black). Bars correspond to the percentage of proteins annotated with each GO term. Asterisks indicate significant enrichment relative to the number of identified proteins (EASE score) * ≤ p=0.05, ** ≤ p=0.005, *** ≤ p=0.0005, **** ≤ p=0.00005, exact p-values are described in Supplemental table 2.

The resistance of *usp18^C61A/C61A^* mice to a plethora of viral infections [63] has raised the possibility of USP18 inhibition as a host-directed antiviral strategy. Our data indicate that enhanced ISGylation can be protective *in vivo* following acute bacterial infection of the lung, so we next sought the molecular underpinnings of this protection by determining if ISG15 substrates from *usp18^C61A/C61A^* mice fell into distinct molecular functions relative to wildtype using Gene Ontology analysis. We compared Cluster 2 (unique to wildtype), Cluster 4 (unique to enhanced ISGylation), pooled ISG15 sites (ISGylated), and ubiquitin sites from Cluster 3 (ubiquitinated) to all identified proteins from the input using gene ontology analysis (Fig 3B and S2 Table). We looked for significantly enriched terms in Biological Process, Molecular Function and Cellular Compartment. Cluster 4 was highly enriched in proteins associated with the following GO groups: immune system process, innate immune response, and response to bacterium. Among Molecular Functions enriched in Cluster 4 were protein binding, nucleotide binding and ATP binding though these groups were also enriched across all ISGylated proteins (Fig 3B and S2 Table). We ranked enriched ISG15-substrates in Cluster 4 based on the total number of sites per protein and focused on substrates with additional sites that were unique to conditions of enhanced ISGylation. ISGylation of these immune system regulators could potentially activate or inhibit their function resulting in better lung capacity in *usp18^C61A/C61A^* animals.

### Host viral restriction factor SAMHD1 is modified by ISG15 following bacterial infection

One of the most ISGylated proteins in Cluster 4 which affects both innate immune responses and cytosolic nucleotide levels is the sterile alpha motif and histidine-aspartate-domain-containing protein 1 (SAMHD1). SAMHD1 is a viral host restriction factor for HIV and other retroviruses which depletes cytosolic dNTPs [64] and can also act as a single-stranded 3’ exonuclease [65]. These activities suppress the innate immune response including the Type I Interferon pathway [66-68]. Indeed, SAMHD1, like ISG15, leads to an interferonopathy in human patients who bear mutations in the gene [69-73]. Here we found SAMHD1 modified by ISG15 on twenty-one distinct sites following *Francisella* infection in the lung (Fig 4A and S3 Table). We also previously identified SAMHD1 as an ISG15 target following *Listeria* infection in the liver but following *Francisella* infection in the lung we mapped many more sites (Fig 4 A-B) [49]. Four of the twenty-one ISG15 target lysine sites are upregulated only in conditions of enhanced ISGylation. When we mapped these ISGylation sites onto the solved crystal structure of the SAMHD1 tetramer [74], it was apparent that ISGylation could indeed directly impinge on cofactor and substrate binding (Fig 4C, highlighted in red). Moreover, SAMHD1 is in a constant equilibrium between monomeric, dimeric, and tetrameric forms with the formation of a tetramer being initiated by the binding of GTP. In addition to hindering GTP binding, the ISGylation sites could sterically hinder the formation of dimers and tetramers, as shown in the modeling of ISG15 on the monomer and dimer of SAMHD1 (Fig 4D, pink arrows) [75, 76]. One possibility is that ISGylation of SAMHD1 could be induced to temper catalytic activity after the resolution of infection and thus we could detect these sites because USP18 is catalytically inactive (*usp18^C61A/C61A^*). Notably, SAMHD1 is modified following both *Listeria* and *Francisella* infection suggesting the potential for a new mode of post-translational regulation of SAMHD1 by ISG15.

**Fig 4.**
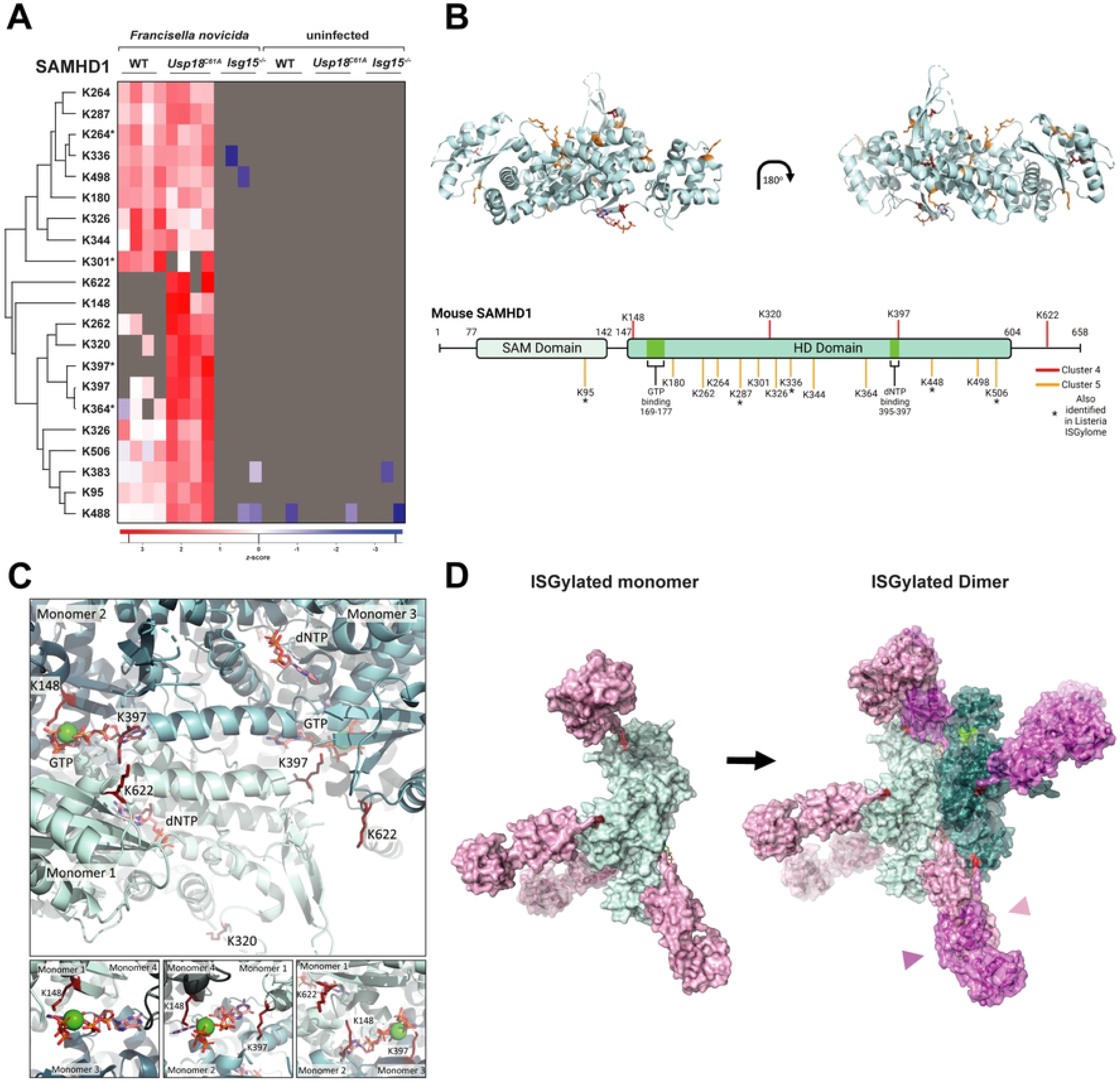
ISG15 modification of the innate immune response controlling protein SAMHD1 may hinder tetramerization and function. (A) Heatmap of non-imputed values generated from non-supervised hierarchical clustering of all SAMHD1 peptides modified by ISG15 identified as significant from the two-way ANOVA following GlyGly(K) enrichment. ^ indicates a peptide that had either methylation-M or carbamylation-C in addition to the GlyGly modification. (B) Modeled SAMHD1 (6brh) with indicated ISGylated sites both in ribbon and cartoon diagram created with biorender.com. (C) Large and zoomed-in view of SAMHD1 (6brk) GTP and dNTP active sites formed from tetramerization, highlighting modified lysines from cluster 4 showing how binding of ISG15 would inhibit the formation of the individual enzymatic pockets. (D) 3D model of four ISG15 molecules bound to SAMHD1 at K148, K320, K397, and K622 the sites identified only in *usp18^C61A/C61A^* samples both as a monomer and dimer.

### Validation of ISGylation on SAMHD1 and potential effects on function

We next sought to validate the covalent modification of ISG15 on SAMHD1. We differentiated human THP-1 wildtype and *samhd1^-/-^* monocytes into macrophages using PMA and infected the cells with *F. novicida* for four hours [77]. We lysed the cells and performed an immunoprecipitation (IP) of endogenous SAMHD1 to confirm SAMHD1 expression in wildtype but not SAMHD1-deficient cells. While undifferentiated SAMHD1-deficient monocytes had increased ISGylation prior to PMA treatment, as reported in the literature [78, 79] in mouse macrophages, there was distinctly less ISGylation than in wildtype cells after 24 and 48 hours of PMA treatment suggesting a lower overall threshold for prolonged ISGylation (Fig S2A). Following SAMHD1 immunoprecipitation (IP) we probed for ISG15 and observed several upper bands in the wildtype cells that are absent in the SAMHD1-deficient cells. The SAMHD1-ISG15 complex appeared to migrate around 90 kDa in denaturing conditions with additional upper bands around 120 kDa and thus would represent a multi-ISGylated SAMHD1 molecule as indicated by our proteomics analysis (Fig 5B, indicated by arrows). Since the slower-migrating ISG15 bands were difficult to resolve, we also validated our IP via LC-MS/MS and were able to identify peptides from ISG15 associated with SAMHD1 in wildtype infected samples which were absent in our *samhd1^-/-^* samples (Fig 5C).

**Fig 5.**
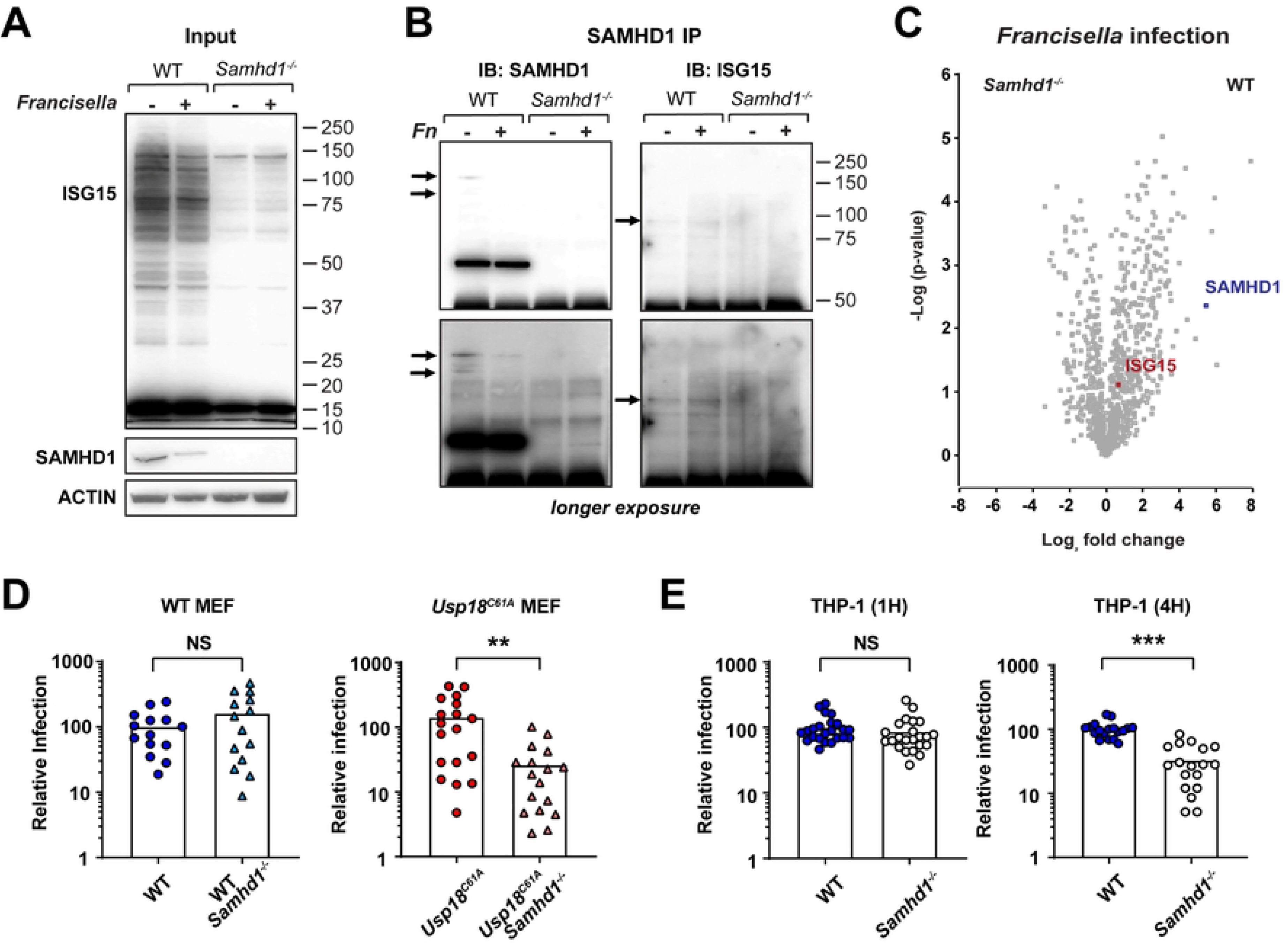
SAMHD1 is modified by ISG15 and reveals a relationship between SAMHD1 and *Francisella* infection. (A-C) Wild-type and *Samhd1^-/-^* THP-1 monocytes were differentiated into macrophages via 100ng/mL PMA stimulation. 1x10^7^ cells were infected with *F. novicida* for four hours, then lysed with Triton (A) lysate was run on SDS-PAGE, blotting for the indicated proteins. (B) Lysate was immunoprecipitated for SAMHD1 and visualized on SDS-PAGE gel for SAMHD1 and ISG15 Arrows indicate upshifted ISG15 bands corresponding to SAMHD1 modification. (C) Lysate was immunoprecipitated for SAMHD1 and run on LC-MS/MS, the log fold change of differentially expressed proteins in infected WT vs *Samhd1^-/-^* samples is shown via volcano plot with SAMHD1 and ISG15 highlighted. (D) CFUs from *Samhd1^+/+^* or CRISPR-Cas9 generated *Samhd1^-/-^* MEFs infected with *F. novicida,* MOI of 100 for 24 hours. Wild-type values were normalized to one hundred; data from three independent experiments with n=4-5/group is shown analyzed Mann-Whitney test, **p=0.0011. (E) Differentiated THP-1 cells were infected with *F. novicida* for indicated time points. 1-hour cells were washed three times with PBS before lysis with 0.1% triton in water and serially diluted and plated for CFUs. Gentamicin was added at 1 hour to the cells infected for 4 hours and they were lysed as before. Statistical significance was determined via Mann-Whitney analysis, adjusted p-values are shown; *** < p=0.0005.

Since our structure-function analysis suggests that ISG15 modification of SAMHD1 in conditions of enhanced ISGylation could reduce SAMHD1 enzymatic activity, we subsequently tested the capacity of bacteria to replicate in cells without SAMHD1. We performed a gentamicin protection assay to quantify intracellular bacterial colony-forming units (CFUs) at 24 hours post-infection. Since fibroblasts are relatively resistant to *Francisella* infection, we used an MOI of 100 of *F. novicida*. There was a significant SAMHD1-dependent decrease in bacterial burden in *usp18^C61A/C61A^* MEFs whereas in wild-type cells, bacterial load did not change upon SAMHD1 deletion (Fig 5D Fig S2B). From these data, we hypothesize that when ISGylation is dysregulated SAMHD1 activity facilitates *F. novicida* replication. We subsequently tested the antibacterial capacity of SAMHD1-deficient THP-1 derived PMA-activated macrophages to clear bacterial infection relative to wildtype [77]. SAMHD1-deficient macrophages also had fewer colony forming units of *Francisella* than wildtype cells (Fig 5E). In both cases the combination of increased ISGylation and SAMHD1 permit bacterial replication, whereas SAMHD1 deletion under these conditions reduces bacteria burden.

To our knowledge, this is the first time that anyone has assessed the role of SAMHD1 in bacterial infection. These data further indicate that bacteria can take advantage of SAMHD1 activity during infection, which correlates with the ability of *Francisella* to exploit innate immune signaling for its own benefit. Finally, our work implicates a new function for ISG15 as a post-translational modification with the capacity to regulate SAMHD1 activity which will be a fruitful avenue to explore in future studies.

## Discussion

Here we explored the role of ISG15 and ISGylation in host responses to the intracellular bacterial pathogen *Francisella novicida*. We found that *Francisella* induces robust ISGylation in macrophages and *in vivo* in the lung, liver, and spleen following intranasal infection. Macrophages with unchecked ISGylation cannot control infection as well as wildtype cells. Following infection *in vivo*, there is not a significant difference in bacterial load, but ISG15-deficient animals have increased lung lesions and dyspnea, whereas mice with enhanced ISGylation have reduced lesions and improved lung function. We mapped the ISGylome in the lung following infection and found increased modification of enzymes that affect innate immune signaling. We validated the ISGylation of SAMHD1 and assessed the role of SAMHD1 on bacterial replication, finding that the lack of SAMHD1 results in reduced bacterial burden. Taken together, our work contributes the first comprehensive ISGylome of lung tissue and identifies that increased ISGylation can protect lung function following bacterial pneumonia (Fig 6).

**Fig 6.**
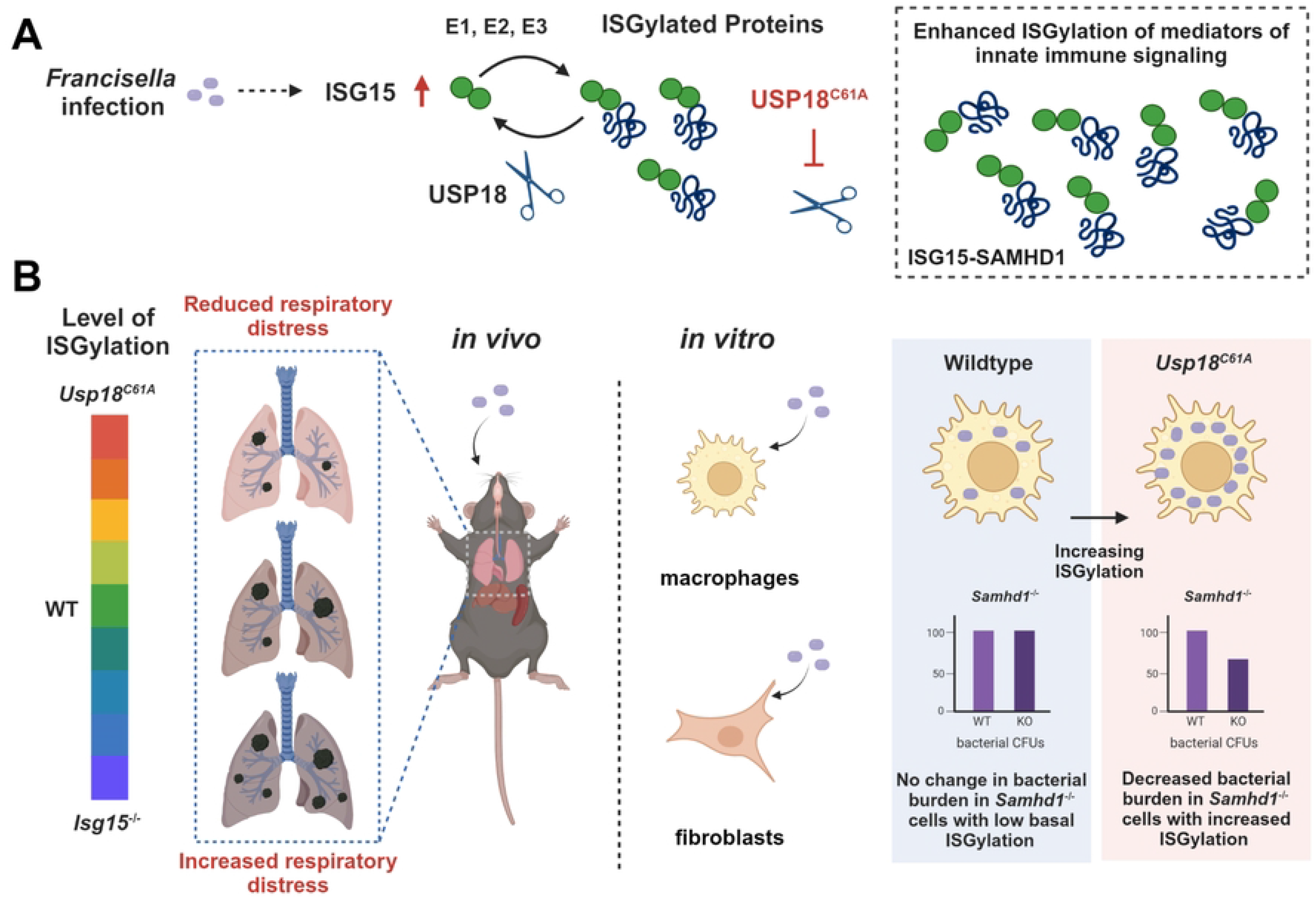
Graphical abstract of the effect of ISG15 during respiratory *F. novicida* infection. (A) Depiction of the increase in ISG15 and ISGylation following *F. novicida* infection, highlighting the process of ISGylation and the inhibition of the removal of ISG15 from proteins in the *Usp18^C61A/C61A^* model. The box represents the enrichment of ISGylated immune system mediators, including SAMHD1, identified via mass spectrometry in the lungs of mice with enhanced ISGylation. (B) Depiction of the ISG15 dependent *in vivo* and *in vitro* responses to *F. novicida. In vivo,* there is a decrease in respiratory distress with increasing levels of ISGylation that is reversed in the absence of ISG15. *In vitro,* enhanced ISGylation leads to increased *F. novicida* burden, and the removal of SAMHD1 from a system with increased ISGylation, but not basal levels of ISGylation, results in decreased bacterial load.

### Lung and Liver ISGylomes Following Acute Bacterial Infection Show Striking Similarity

To understand the mechanism of action of ISG15 several studies have employed proteomics to identify the ISGylome of all ISG15-modified proteins following interferon treatment. Most early reports were unable to map the lysine sites that are modified by ISG15, since ISG15 and ubiquitin bear the same C-terminal LRLRGG sequence [80-82]. Recently, we pioneered a genetic approach to distinguish ISG15 modification sites from ubiquitin and mapped the ISGylome of the liver from *Listeria-*infected mice [49]. Here, we used this method to identify the ISGylome of lung tissue from *Francisella novicida-*infected mice. To our knowledge, this study is the first to map the lung ISGylome following a bacterial or viral respiratory infection. Compared to the *Listeria-*induced liver ISGylome, we identified fewer ubiquitin sites prior to infection (Cluster 1, 107 sites on 75 proteins) however many ubiquitin sites were induced by infection (Cluster 3, 831 sites on 404 proteins) rather than lost. ISGylation in the lung is induced in three distinct clusters: one which is common to wildtype and *usp18^C61A/C61A^* mice (Cluster 5), and two others which are unique to wildtype (Cluster 2) and *usp18^C61A/C61A^* (Cluster 4), respectively. The pattern is like that observed in the liver but relatively there are half the amount of ISG15 sites in the lung (515 unique sites) compared to those in the liver (929 sites). Notably, 91 of these sites occur on identical lysines and were identified as common following *Francisella* and *Listeria* infection. Since sites are conserved down to the lysine residue both in liver and lung, this could indicate a core list of site-specific ISG15 targets that are common to intracellular bacterial infection. For ubiquitin and other ubiquitin-like proteins target specificity is determined by E3 ligases [83] or a consensus sequence [84]. Since there is not a discernable consensus sequence for ISG15 it is tempting to speculate that there could be additional ISG15 E3-ligases, distinct from HERC5, induced following bacterial infection. Recent structural studies may elucidate whether there is an E2 specific-binding surface that could confer specificity [85, 86].

### ISGylation of SAMHD1: A New Mechanism of Post-Translational Control

Gene ontology analysis of ISGylated targets in the *usp18^C61A/C61A^* mouse lungs revealed nearly ∼15% enrichment in the GO Biological Process terms: innate immune response, immune system process, and 10% enrichment in response to bacterium relative to a 2-4% baseline enrichment of proteins associated with those terms between in the lung. Interestingly, the GO Molecular Function terms had nearly ∼ 30% enrichment of GO terms involving nucleotide binding and ATP-binding, which was a common feature specific to ISGylated targets rather than ubiquitin substrates in the lung. Following *Listeria monocytogenes* infection in the liver, these GO terms were also enriched for ISG15-substrates [49]. For validation and further characterization, we chose to focus on SAMHD1 as a covalent substrate of ISG15, because it met the criteria of being enriched in both GO groups: innate immune function and nucleotide binding. Interestingly, SAMHD1 is differentially modified in wildtype and *usp18^C61A/C61A^* animals following *Francisella* infection. ISG15-deficient patients also exhibit an Aicardi-Goutières-like interferonopathy [87], which phenocopies some patients with mutations in SAMHD1, who also have increased Interferon signaling in the brain [69-73].

SAMHD1 has primarily been studied during retroviral infection, where it restricts viral replication by depleting host dNTPs [64, 88, 89]. This depletion of dNTPs also functions to limit cGAS/STING signaling and Type I interferon production in HIV infected cells [78, 79, 90]. Since bacteria do not pirate dNTPs from the host for their own growth and replication like viruses do, SAMHD1 activity has not yet been explored in the context of intracellular bacterial infection. Lipopolysaccharide (LPS) priming has been reported to lead to SAMHD1-mediated suppression of NF-kB and Type I Interferon signaling [90, 91]. We have shown here that SAMHD1 promotes *Francisella novicida* replication in THP-1-derived human macrophages and *usp18^C61A/C61A^* mouse embryonic fibroblasts, both of which exhibit increased basal ISGylation. Macrophages exhibit this due to PMA treatment, while fibroblasts do so due to a USP18 mutation. Conversely, SAMHD1 deletion in wildtype fibroblasts did not affect bacterial load, suggesting that SAMHD1 confers a pro-bacterial effect in this cell type only under conditions of enhanced ISGylation. Recent work on other mediators of innate immune signaling implicated ISGylation as a post-translational modification required for higher order oligomerization of complexes such as MDA5 and RNF213 [92, 93]. Indeed, Gack and colleagues found that ISGylation is required for MDA5-driven signaling and SARS-CoV-2 encodes a protease which can remove ISG15 from MDA5. The sheer number of ISG15 sites identified on SAMHD1 makes it compelling to hypothesize that ISG15 may also affect higher order complexes of SAMHD1. Furthermore, the identification of specific sites solely when USP18 is inactive could implicate the enzyme as a regulator of ISG15-mediated control of SAMHD1. Ultimately, our unbiased proteomics approach revealed a new post-translational modification of SAMHD1 conserved in the liver and lung, which can impinge on intracellular bacterial replication. Future studies will address how ISGylation affects SAMHD1 structurally and whether SAMHD1-deficiency in mice affects systemic bacterial infection *in vivo*.

### The role of ISG15 regulation in clearance of viral and bacterial respiratory infections

ISG15 plays a critical role in lung host defense, which was brought to light by the discovery that human patients who lack ISG15 expression are susceptible to the BCG vaccine [34]. In these patients, Bogunovic and colleagues pinpointed the role of ISG15 as a cytokine to propagate Interferon-γ production. More recently, patients with pneumonia of unknown etiology were diagnosed with loss of function mutations in ISG15, highlighting the importance of ISG15 and ISGylation for proper lung function [94]. In general, covalent modification by ISG15 or ISGylation is a protective host response against both viral and bacterial respiratory infections, such as Sendai virus, Respiratory Syncytial Virus (RSV), and infection with *Mycobacterial* species [33, 45, 47, 95]. During Sendai virus infection, mice deficient in either ISG15 or its E1 enzyme, UBE1L (required for ISGylation), have increased mortality relative to wildtype mice. *Ube1L^-/-^* mice also have fewer inflammatory cells recruited to the lung than either wildtype or *isg15^-/-^* mice, indicating a unique role for ISGylation in immune cell recruitment and the resulting airway obstruction [96]. These data mirror our own observations following *Francisella* infection as indicated by whole-body plethysmography and histology of mouse lungs following infection. In addition, mouse survival during influenza A viral infection was ISG15-conjugation dependent but was not the result of differences in viral load [96]. Similarly, in response to *Mycobacterium tuberculosis* infection, the lack of ISGylation did not significantly affect bacterial burden however it did reduce mouse survival indicating ISGylation-dependent mechanisms responsible for aberrant pathology [47]. Our study contributes similar findings for *Francisella* infection in that bacterial colony-forming units remain the same with enhanced or absent ISGylation, but the cellular infiltrates (pyogranulomas) are reduced with corresponding improved airway lung function. Since *F. novicida* is such a potent mouse pathogen, survival under conditions of enhanced ISGylation was only extended by hours (data not shown). Yet for both *Mtb* and *Francisella* infection, ISGylation protects from severe lung damage.

Antibiotic resistance and the ongoing threat of emerging viral infections has led to a renewed interest in host-directed antiviral and antibacterial therapies [97]. While the lack of ISGylation through UBE1L deletion can differentiate between the role of free and conjugated ISG15, *usp18^C61A/C61A^* mice more directly model the consequences of USP18 inhibition [63]. Other groups have used these mice to describe the protective nature of de-ISGylation during VACV and Influenza B infections, wherein the mutation of USP18 results in drastically reduced infection compared to total ISG15 knock out alone [63]. In these cases, the regulation of the ISG15 conjugation system has proved to impact disease differently than the deletion of ISG15 alone. Our study adds to these findings by indicating that the inability to remove ISG15 from target proteins reduces respiratory distress following acute *Francisella* infection, whereas the removal of ISG15 entirely leads to exacerbated lung pathology. This result suggests that inhibition of the deconjugase activity of USP18 could lessen pneumonic-like symptoms in both bacterial and viral infections. Other isopeptidases specific to ubiquitin have been targeted in the context of cancer [98, 99]. The pronounced antipathogenic effect following viral infection [63] combined with the protection from mortality [78] and improved respiratory function that we observed following acute bacterial infection justify exploring prophylactic or post-diagnosis USP18 inhibition for treatment and prevention of respiratory infections.

## Acknowledgements

We thank the Radoshevich laboratory for comments on the manuscript. We thank Dr. Lee-Ann Allen for her generous gift of *F. novicida* (U112) and many insightful conversations. We thank Dr. Brad Jones, and Dr. Mike Apicella for imparting valuable knowledge about *Francisella* to us. We thank Dr. Steve Varga and Stacey Hartwig for helpful discussions and use of their Buxco apparatus. We thank Dr. Li Wu for helpful discussions and providing us with SAMHD1-deleted THP-1 cells. The pSpCas9(BB)-2A-GFP (PX458) plasmid was a gift from Feng Zhang (Addgene plasmid #48138; http://n2t.net/addgene:48138; RRID: Addgene_48138). We used the cell sorter to obtain CRISPR/Cas9 clones at the Flow Cytometry Facility, which is a Carver College of Medicine/Holden Comprehensive Cancer Center core research facility at the University of Iowa. The facility is funded through user fees and the generous financial support of the Carver College of Medicine, Holden Comprehensive Cancer Center, and Iowa City Veteran’s Administration Medical Center. The LC-MS/MS samples were run by Dr. Anthony Saviola at the University of Colorado, Anschutz, Mass Spectrometry Proteomics Shared Resource Facility (RRID: SCR_021988) which is supported by the Cancer Center Support Grant (P30CA046934).

## Materials and methods

### Ethics Statement

All mouse experiments were approved by the Institutional Animal Care and Use Committee at the University of Iowa under protocol 2032090.

### Materials

For immunoblotting, we used anti-ISG15 antibodies from Santa Cruz (F-9, sc-166755) at 1:200, and Sino-biological (12729-R239) at 1:2000. We used an anti-β actin antibody from ThermoFisher (MA1-140) at 1:5000. For both immunoblotting and immunoprecipitation, we used a polyclonal anti-SAMHD1 antibody from Abcam (ab67820) at 1:1000 or 1µg/sample respectively. For immunoprecipitation, we used Pierce^TM^ Protein A/G Magnetic Agarose Beads from ThermoFisher (78609).

### Mammalian cell growth conditions

All cells were cultured in a sterile environment and grown at 37°C with 5% CO_2_. Mouse Embryonic fibroblasts were grown in DMEM with Glutamax (Gibco, Waltham, MA) supplemented with 10% Fetal bovine serum. Bone marrow-derived macrophages were obtained via dislocation of mouse femur and tibia and placed in sterile un-supplemented RPMI 1640 media (ThermoFisher 11875093) on ice. Bones were washed with ethanol and RPMI following which, one end was cut and placed into a PCR tube which was then placed in a 1.5ml Eppendorf tube containing 100µl of RPMI media. PCR/Eppendorf tube sets were spun down at 8000xg for 30sec. Pellets were then resuspended and transferred to new tubes which were then spun down at 300xg for 5min at 4°C and red blood cells were lysed with 0.83% NH_4_Cl. The remaining cells are plated on uncoated 10cm dishes at 3.5x10^6 cells/plate. Cells are incubated for 7 days in RPMI supplemented with 10% fetal bovine serum, 1% glutaMAX (Life Technologies, 35050061), 1% 1M HEPES (Life Technologies, 15630080), 1% Sodium pyruvate (Life Technologies, 11360070), 2-Mercaptoethanol at 0.05M (VWR, 97064-878), and 20% cultured media from L929 cells: NCTC clone 929 [L cell, L-929, derivative of Strain L] (ATCC CCL-1) for differentiation into macrophages. Following differentiation cells were lifted with accutase (Sigma-Aldrich A6964) and either frozen or plated for further experiments. THP-1 cells both WT and *samhd1^-/-^* were a gift from the Wu lab at the University of Iowa. They were cultured with RPMI media supplemented with 10% fetal bovine serum, 0.05M 2-Mercaptoethanol, and 1µg/ml of puromycin. They were differentiated into macrophages by the addition of 100ng/ml PMA for 24hrs.

### Development of CRISPR/Cas9 cell lines

The SAMHD1 knockout fibroblast cell lines were generated by using a CRISPR/Cas9 approach. Target sequences were designed with Benchling (benchling.com). Oligonucleotides were synthesized by IDT (Coralville, IA, USA) and cloned into the pSpCas9(BB)-2A-GFP (PX458) plasmid which was a gift from Feng Zhang (Addgene plasmid #48138; http://n2t.net/addgene:48138; RRID: Addgene_48138). Cells were transfected with the SAMHD1-targeting plasmids with FuGene HD. 48 h post-transfection, GFP-positive cells were sorted and plated in a 96-well plate for single-clone selection with Becton Dickinson FACS Aria Fusion (BD Science). SAMHD1 deficiency was tested by Interferon-α stimulation and subsequent western blotting for SAMHD1. The absence of SAMHD1 protein expression was confirmed by immunoblotting as well as genomic PCR and next-generation sequencing of the targeted gene segment (PCR sequence alignment is represented in Supplemental Table 4).

### Bacterial growth conditions and infections

*Francisella novicida* (U112) a gift from the Allen Lab at the University of Missouri, was grown on Muller Hinton agar plates (prepared as described in Conlan et al 2011). Overnight cultures were grown in Muller Hinton broth. The absorbance of the culture was read and an OD between 0.4-0.7 was used for all experiments. CFUs/mL were calculated with an OD of 1= 5x10^8 CFUs/mL. Bacteria were then washed three times with 1xPBS and resuspended in serum-free media. Bacteria were added to serum-free mammalian cell culture media at the indicated MOI. A fixed volume was then added to each well of cells in 24-well plates. The plates were then spun down at 800 rpm for 1 min to synchronize infection. Cells were incubated with the bacteria for 1hr at 37°C, with 5% CO_2_. Serum-free media was then replaced with complete media containing 20µg/ml of gentamicin (Thermo Fisher 15750060) to kill extracellular bacteria. The cells were washed with room temperature 1xPBS and then harvested at the indicated time points by lysis with 0.1% triton in H_2_O and serially diluted in PBS before plating onto Muller Hinton plates, from which colony forming units (CFUs) were enumerated.

### *In vivo* infections

A single colony of *Francisella novicida* was grown up in 5mL Muller Hinton broth overnight. The overnight culture was then subcultured into 50mL at a 1:10 dilution and grew to OD between 0.4-0.7 mid to late log phase. Bacteria were aliquoted and snap frozen in 100µL volumes. Serial dilutions were plated out to determine the number of CFUs per aliquot. C57BL/6 mice (*Isg15^+/+^*, *Isg15^-/-^*, and *Usp18^C61A/C61A^*) between 8-12 weeks of age were infected intranasally following isoflurane immobilization with 100 CFUs per animal. Mice were evaluated daily for weight loss and, in some cases, pulmonary function using a whole-body plethysmograph (Data Sciences International, New Brighton, MN) to measure changes in respiration from baseline measurements taken prior to infection. Enhanced pause (Penh) and mid tidal respiratory rate (f) parameters were calculated based on pressure and volume changes in the chamber caused by respiration and averaged over a 5-min period. Mice were euthanized on day three post-infection via cervical dislocation. Tissues were removed and homogenized in one mL of sterile PBS with a PowerGen 125 homogenizer (Marshall Scientific, FS-PG125). For assessment of CFUs, Homogenate was serial diluted in PBS and plated on Muller Hinton plates. For the SDS-PAGE and Mass-Spectrometry analysis of animal samples, tissue homogenates were centrifuged at 3435 x g for 10 min at 4°C the soluble fraction below the layer of fat was removed and resuspended in a 1:1 ratio with 10% SDS. The concentration of protein was then assessed via BCA and samples were normalized for downstream application. For histological analysis, organs were extracted from mice and placed in 10% NBF formaldehyde. Tissues were then processed (i.e., routinely dehydrated through a progressive series of alcohol and xylene baths), paraffin-embedded, sectioned (∼4 µm) and stained by hematoxylin and eosin (HE) by the Comparative Pathology Laboratory (University of Iowa). Tissue were examined and imaged by a boarded veterinary pathologist (DKM) using the post-examination method of group masking [100].

### SDS-PAGE

For the SDS-PAGE analysis, cells were washed with 1x PBS then lysed in 1% Triton lysis buffer or 1x RIPA lysis buffer supplemented with Complete Mini Protease inhibitor cocktail (Roche). The concentration of protein was then assessed via BCA and samples were normalized and added to sample buffer supplemented with 10% BME before being run on an SDS-PAGE gel (Invitrogen NW04125BOX). Gels were transferred using an iBLOT transfer system (Invitrogen), blocked in 5% milk for 1 hour at RT, incubated with primary antibody overnight at 4 °C, washed with 0.05% Tween in PBS three times (each wash for 7 min), and incubated for 1 h at room temperature with secondary antibody coupled to HRP. Blots were washed again three times with 0.05% Tween in PBS and revealed using ECL Western Blotting Substrate (Pierce, Waltham, MA).

### Immunoprecipitation

Cells were lysed and protein concentration was determined with a BCA assay. 500µg of lysate was incubated with 1µg of anti-samhd1 antibody (ab67820) overnight rotating at 4°C. The sample was then incubated for ∼4 hours with magnetic A/G beads (Thermo Scientific 78609) again rotating at 4°C. The beads were pulled down and washed 3x with cold PBS and the antibody/conjugate was eluted off the beads by the addition of Sample buffer with 10% BME and boiled for 5min, mixing every min. The elution was then run on an SDS-PAGE Gel as described above. For Mass-spectrometry analysis samples were cleaved with trypsin to elute from the magnetic A/G beads.

### Proteomics sample preparation

The protein concentration from animal organ lysates was measured via BCA assay (Pierce) and equal protein amounts containing 2mg total protein each were used for further analysis. Each sample was digested in a Suspension Trapping (S-trap) midi spin column (Protifi, USA) according to the manufacturer’s instructions. Proteins were reduced by adding 4.5mM (final concentration) dithiothreitol and incubated for 30min at 55⁰C. Proteins were then alkylated with 10mM (final concentration) iodoacetamide for in the dark for 15min at room temperature. Samples were then acidified by the addition of phosphoric acid to a 1.2% final concentration. The samples were then mixed with 6x volume of a binding buffer (90% methanol; 100mM (final concentration) TEAB) and loaded onto an S-trap filter and centrifuged at 3665xg for 30s. The columns were then washed three times with a wash solution (90% methanol; 100mM (final concentration) TEAB) before digestion with 100ug trypsin (Promega) (1/10, w/w) overnight at 37⁰C.

The S-trap filters containing digested peptides were then rehydrated for 30min at room temperature with 500ul of elution buffer 1 (50mM TEAB (final concentration) in water). Subsequent elution steps of 500ul elution buffer 1 (0.5% Trifluoracetic acid in water) followed by 500ul elution buffer 2 (50% Acetonitrile in water) completed the peptide elution. The three subsequent eluates containing peptides were pooled. At this point, 100ug of digested material was aliquoted and subsequently de-salted with reverse-phase C18 OMIX tips (Pierce), all according to the manufacturer’s specifications before proceeding to LC-MS/MS for Shotgun proteomics analysis. The remaining peptides were then lyophilized for 48 hours. Immunocapture of di-gly peptides was performed using Anti-diglycine lysine antibody conjugated agarose beads (PTM Bio) according to the manufacturer’s instructions. Lyophilized peptides were reconstituted in 200ul immunoprecipitation buffer (100nM NaCl, 1mM EDTA, 20mM Tris-HCL (final concentrations) pH 8.0). Peptides were incubated with anti-diglycine lysine conjugated agarose overnight at 4⁰C on a rotator. The agarose was washed three times in wash buffer (100mM NaCl, 1mM EDTA, 20mM Tris-HCL (all final concentrations) pH 8.0) followed by two washes in Milli-Q grade water. Modified peptides were eluted three subsequent times in 0.1% trifluoracetic acid and desalted on reverse-phase C18 OMIX tips (Pierce), all according to the manufacturer’s specifications. Purified di-gly modified peptides were dried under vacuum and stored at -20⁰C until LC-MS/MS analysis.

### LC-MS/MS and data analysis

Digested peptides were loaded onto individual Evotips following the manufacturers protocol and separated on an Evosep One chromatography system (Evosep, Odense, Denmark) using a Pepsep column, (150 um inter diameter, 15 cm) packed with ReproSil C18 1.9 um, 120A resin. The system was coupled to the timsTOF Pro mass spectrometer (Bruker Daltonics, Bremen, Germany) via the nano-electrospray ion source (Captive Spray, Bruker Daltonics). The mass spectrometer was operated in PASEF mode. The ramp time was set to 100 ms and 10 PASEF MS/MS scans per topN acquisition cycle were acquired. MS and MS/MS spectra were recorded from m/z 100 to 1700. The ion mobility was scanned from 0.7 to 1.50 Vs/cm2. Precursors for data-dependent acquisition were isolated within ± 1 Th and fragmented with an ion mobility-dependent collision energy, which was linearly increased from 20 to 59 eV in positive mode. Low-abundance precursor ions with an intensity above a threshold of 500 counts but below a target value of 20000 counts were repeatedly scheduled and otherwise dynamically excluded for 0.4 min.

Data analysis was performed with Fragpipe (version 17.1) using the MSFragger (version 3.4) search engine configured with Philosopher (version 4.2.1) with default search settings for LFQ-MBR workflow including a false discovery rate set at 1% for both the protein and peptide level. Two separate searches were performed to analyze spectra generated from the input shotgun as well as di-gly enriched samples. In both searches, spectra were searched again mouse proteins in the Uniprot database (Taxonomy ID 10090, 55,315 protein sequences) as well as *Francisella tularensis subsp. novicida* str. U112 (Taxonomy ID 401614, containing 1,719 protein sequences). For both searches, a mass tolerance for precursor ions was set at a range of -20 to 20ppm with a mass tolerance for fragment ions at 20ppm. Enzyme specificity was set to trypsin and for the input shotgun search, a maximum of two missed cleavages was set, while the di-gly enriched samples were set to an allowance of three missed cleavages. Carbamidomethylation of cysteine residues was set as a fixed modification with variable modifications set to oxidation of methionine and acetylation of protein N-termini. For the di-gly enriched samples, GlyGly modification of lysine residues was added as an additional variable modification. In both searches, match between runs was enabled with retention time tolerance window set to 1 minute and an ion mobility tolerance set to 0.05 (1/K_0_). Only proteins with at least one unique or razor peptide were retained in the shotgun search.

To further analyze di-gly sites, the combined modified peptide table generated from MSFragger was loaded into Perseus software (version 1.6.14.0). Potential contaminants were removed. Remaining hits were considered as identifications. Intensities were log2 transformed and normalized for each sample by subtracting the median intensity. Replicate samples were grouped, and the identifications with less than three valid values in at least one group were removed, deeming a hit to be quantifiable. Missing values were imputed from a normal distribution around the detection limit. For statistical analysis, replicate sample groups were defined based on genotype: WT, ISG15^-/-^, USP18^C61A/C61A^ and infection status: uninfected, *F. novicida*. A two-way ANOVA was performed to compare intensities in both the genotype and infection conditions. This test produced a p-value for infection as well as genotype (-log p-value). Sites with a p-value for either condition of less than 0.01 were considered significant. Results of the ANOVA are reported as a heatmap in Figure 3a after non-supervised hierarchical clustering. The cluster analysis of the significantly regulated sites is reported in Supplementary Table 1. For the analysis of the input shotgun data, the combined protein table generated from MSFragger was loaded into Perseus. Potential contaminants were removed. Remaining hits were considered as identifications. Intensities were log2 transformed and normalized for each sample by subtracting the median intensity. Replicate samples were grouped, and the identifications with less than three valid values in at least one group were removed, deeming a hit to be quantifiable. Missing values were imputed from a normal distribution around the detection limit. Quantified input proteins were used for further Gene Ontology analysis. GO terms enrichment analyses were performed using Database for Annotation, Visualization, and Integrated Discovery (DAVID) bioinformatics resources (Huang et al 2009, Sherman et al 2021) Significance was determined by the modified Fishers exact test (EASE score) generated by the program.

## Data availability

The LC-MS/MS data have been deposited to the ProteomeXchange via the PRIDE partner repository with the dataset identifier PDX042443. The Project name is “Identification of the lung ISGylome following intranasal *Francisella novicida* infection” For reviewer access, the account username is and the password is “Od4lil0Z”. All other data are available from the corresponding author upon request.

## Statistical analysis

Tests for normality and lognormality were run to determine Gaussian distribution. For non-parametric distributions, a Kruskal-Wallis test was run and comparisons were made via Dunn’s test post hoc. For parametric distributions, a one-way ANOVA was run followed by Tukey’s test post hoc. For both tests, adjusted p-values are shown. The statistical methods for proteomics analysis and gene ontology are discussed in the proteomics methods.

## Supplemental Information Captions

**S1 Fig. Secondary organ ISG15 induction and *in vivo* burden following *Francisella novicida* infection.**

(A-C) Mice were infected with 100 CFUs of *Francisella novicida* intranasally and monitored for 48 hours after which they were sacrificed, and organs were homogenized in PBS. (A and B) SDS-PAGE of the liver (A) and spleen (B) homogenate probed for ISG15 and actin; *’s represent background banding. (C) CFUs were enumerated from serial dilutions of infected organs at 48 hours; compiled data from three independent experiments n=4-9 mice/group are shown. (D) CFUs enumerated from spleen homogenate 72 hours post-infection; compiled data from two independent experiments n=4-9 mice/group are shown.

**S2 Fig. Characterization of *Samhd1^-/-^* cells following activation treatment.**

(A) WT and *Samhd1^-/-^* THP-1 cells were either untreated or treated with PMA for 24 or 48 hours. Cell lysate was then run on an SDS-PAGE gel (B) SDS-PAGE analysis of lysate from interferon-treated WT and *Usp18^C61A/C61A^* MEFs that were deficient in *Samhd1* following CRISPR/Cas9 genomic editing.

**S1 Table. List of GlyGly(K) sites identified via LC-MS/MS.**

This list contains the GlyGly(K) enriched sites ordered by cluster ranking according to the heatmap shown in Figure 3. Columns from left to right contain the rank number, the cluster number, the UniProt accession number, the gene name, the protein name, the position of the modified lysine residue in the protein sequence, any secondary modified lysine residues, the amino acid sequence of the identified modified peptide, the assigned modifications on the peptide, previously identified modifications, the (-)Log two-way ANOVA p-value for genotype, (-)Log two-way ANOVA p-value for infection, and (-)Log two-way ANOVA p-value for interaction.

**S2 Table. Gene ontology analysis of GlyGly enriched proteins.**

This table contains results from the GO analysis of Ubiquitinated proteins (Cluster 3), ISGylated proteins (Cluster 2,4, and 5), and the individual Cluster 2 and Cluster 5 enhanced in wild-type or usp18C61A/C61A samples respectively were assessed relative to all proteins identified in the lung samples (identified) and all mouse proteins in the Uniprot/Swiss-Prot database. Top GO terms were determined, and significant enrichment p-value (EASE score) was assessed relative to the number of mouse or identified proteins using Database for Annotation, Visualization, and Integrated Discovery (DAVID) bioinformatics resources. Data is separated into sheets for each GO term assessed; Biological process, Cellular compartment, and molecular function.

**S3 Table. List of GlyGly(K) sites on SAMHD1 identified via LC-MS/MS.**

This list contains all the GlyGly(K) modified sites on SAMHD1 as shown in the heatmap from Figure 4A. Columns from left to right contain the cluster number obtained from Table 1, the starting position of the peptide, the ending position of the peptide, the peptide length, the charges on the peptide, the amino acid sequence of the identified modified peptide, the assigned modifications on the peptide, the (-)Log two-way ANOVA p-value for genotype, (-)Log two-way ANOVA p-value for infection, and (-)Log two-way ANOVA p-value for interaction.

**S4 Table. Sequence alignment for SAMHD1 knock-out MEFs.**

This list contains the sequence alignments for genomic DNA isolated from both WT and *Usp18^C61A/C61A^* MEFs following the CRISPR/Cas9 deletion of *Samhd1*.

